# Germline variation contributes to false negatives in CRISPR-based experiments with varying burden across ancestries

**DOI:** 10.1101/2022.11.18.517155

**Authors:** Sean A. Misek, Aaron Fultineer, Jeremie Kalfon, Javad Noorbakhsh, Isabella Boyle, Joshua Dempster, Lia Petronio, Katherine Huang, Alham Saadat, Thomas Green, Adam Brown, John G. Doench, David Root, James McFarland, Rameen Beroukhim, Jesse S. Boehm

## Abstract

Reducing disparities is critical to promote equity of access to precision treatments for all patients with cancer. While socioenvironmental factors are a major driver behind such disparities, biological differences also are likely to contribute. The prioritization of cancer drug targets is foundational for drug discovery, yet whether ancestry-related signals in target discovery pipelines exist has not been systematically explored due to the absence of data at the appropriate scale. Here, we analyzed data from 611 genome-scale CRISPR/Cas9 viability experiments in human cell line models as part of the Cancer Dependency Map to identify ancestry-associated genetic dependencies. Surprisingly, we found that most putative associations between ancestry and dependency arose from artifacts related to germline variants that are present at different frequencies across ancestry groups. In 2-5% of genes profiled in each cellular model, germline variants in sgRNA targeting sequences likely reduced cutting by the CRISPR/Cas9 nuclease. Unfortunately, this bias disproportionately affected cell models derived from individuals of recent African descent because their genomes tended to diverge more from the consensus genome typically used for CRISPR/Cas9 guide design. To help the scientific community begin to resolve this source of bias, we report three complementary methods for ancestry-agnostic CRISPR experiments. This report adds to a growing body of literature describing ways in which ancestry bias impacts cancer research in underappreciated ways.

## Main Text

Cancer outcomes differ widely across individuals of different ancestry groups, due to both socioenvironmental factors^1,2^ and molecular differences in cancer makeup^3,4^. One source of these outcome disparities may be due to differences in treatment response resulting from genetic ancestry-associated germline variants. While much attention has focused on pharmacogenetic associations in this regard, a systematic evaluation of whether relationships between predictive biomarkers and cancer targets may be influenced by ancestry has been challenging due to the absence of reference data. However, recent advances in large-scale profiling of cancer dependencies across hundreds of cellular models derived from patients of varying ancestry now afford the opportunity to systematically evaluate whether ancestry-associated molecular differences translate into differences in cancer dependencies that might affect therapeutic response.

We therefore analyzed ancestry-associated cancer dependencies using data from the Cancer Dependency Map **[Figure 1A]**. This project includes data from genome-scale CRISPR/Cas9 gene essentiality screens across 1070 cancer cell lines reflecting 31 cancer lineages to detect essential genes and their relationships with predictive molecular biomarkers. These data have led to the discovery of multiple dependency-associated somatic alterations including WRN dependence in cell lines with microsatellite instability^5^ and PRMT5 dependence in cells with genomic MTAP deletions^6^ amongst others. Additionally, a similar approach has analyzed ancestry-associated compound sensitivity profiles^7^. We therefore reasoned that systematically analyzing the relationship between computationally inferred cell line ancestry and gene dependency profiles might allow us to identify ancestry-associated dependencies and discover new opportunities for target discovery within and across particular ancestry groups as well as to evaluate whether previously unaccounted for sources of bias may exist.

**Figure 1:**
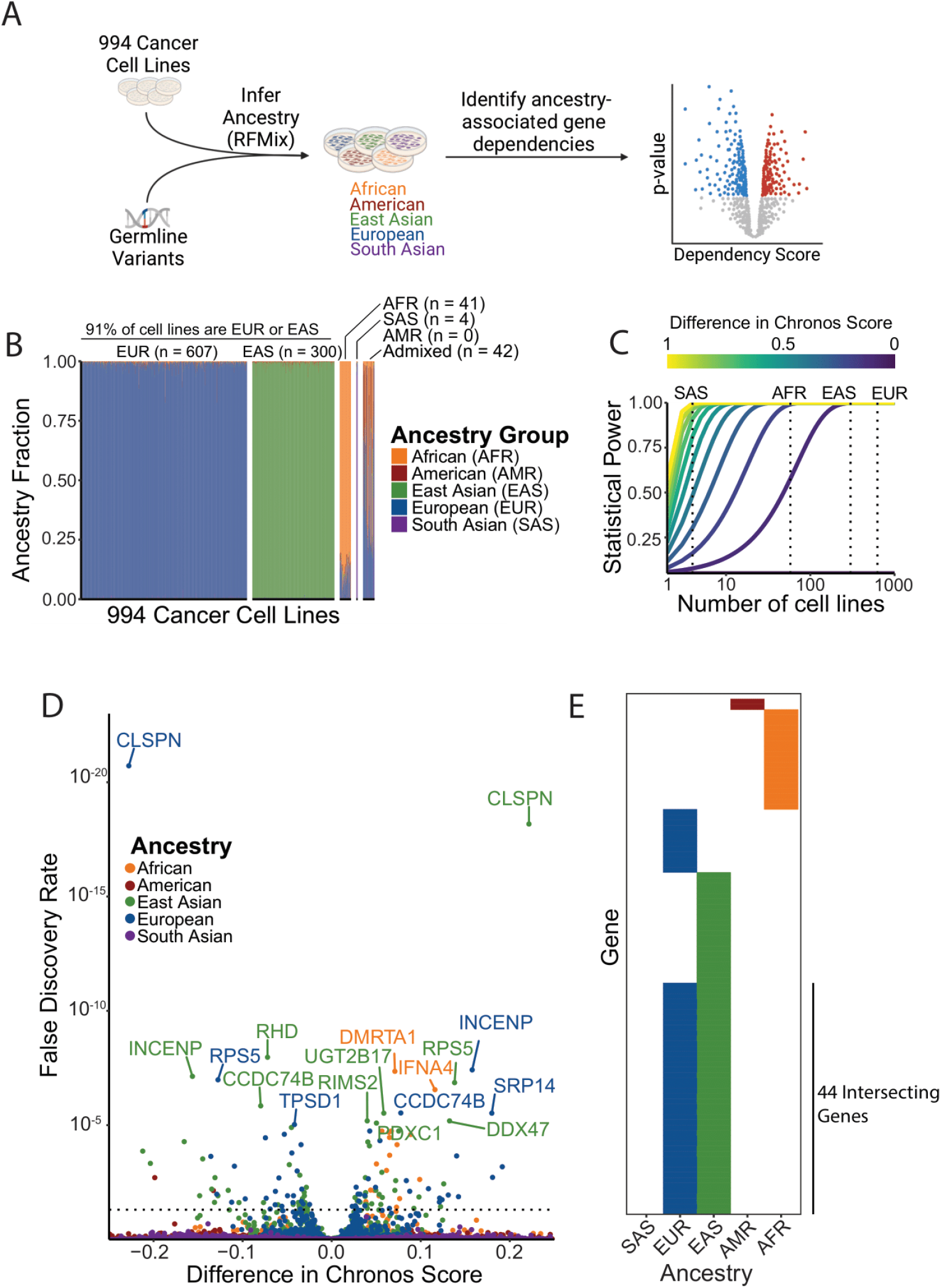
A pan-cancer analysis of ancestry-associated genetic dependencies. **A)** Schematic detailing how the Cancer Dependency Map dataset was leveraged to identify ancestry-associated genetic dependencies. **B)** Genomic fraction for each of five major ancestry groups for 994 the Cancer Dependency Map cancer cell lines. **C)** Power calculation to determine the minimum signal size (difference in Chronos scores) capable of being detected for each ancestry group. A Chronos score difference of 1 represents the difference between “common essential” and unexpressed genes in the Dependency Map. **D)** Volcano plot of ancestry associations among genetic dependencies. **E)** Heatmap indicating the breakdown of ancestry-associated dependencies across ancestry groups.

We systematically cataloged local ancestral haplotypes across the genomes of commonly used cancer cell lines, focusing on the 994 (out of 1829 total) cell line models in the Cancer Cell Line Encyclopedia collection for which publicly available Affymetrix SNP6 germline variant data have been analyzed^8^. While previous reports have evaluated cell line genetic ancestry at genome scale^8–12^, we hypothesized that such global assessments may preclude the discovery of regional germline associations with dependencies. We therefore leveraged germline variants from 10,345,968 SNPs genome-wide to infer local ancestry. Specifically, we divided the genome into blocks comprising 0.2 centimorgans (with a median of 580 SNPs per block) and characterized each block as deriving from one (homozygous) or two (heterozygous) of five major continental genetic ancestry groups: African (AFR), American (AMR), East Asian (EAS), European (EUR), and South Asian (SAS) **[Figure 1B]**. In admixed individuals, individual blocks might derive from two of these ancestries, reflecting both maternal and paternal contributions.

At a global level, our results support previous observations^8,12^ that existing cell lines are overwhelmingly derived from individuals of either EUR or EAS ancestry. We assigned a predominant ancestry to cell lines that derived over 80% of their DNA from that ancestry group and called those without a predominant ancestry “Admixed’’. Of the 994 the Cancer Dependency Map cell lines profiled in this study, over 90% of them are predominantly EUR (61%) or EAS (30%) **[Figure 1B]**. Only 41 (4%) of the cell lines were predominantly AFR. When taking local ancestry into account, the underrepresentation of AFR genetic ancestry was even starker. Cell lines characterized as AFR had large contributions from other (primarily European) ancestries; the average AFR genetic ancestry fraction for AFR cell lines was only 89%. In contrast, the average EUR and EAS ancestry fractions for EUR and EAS cell lines are 98.5% and 98.9%, respectively. Only four cell lines in this analysis were primarily SAS. No cell lines had greater than 80% AMR genetic ancestry, though AMR ancestry did comprise 16.3% of the genomes of Admixed cell lines. These imbalances in cell line ancestry limited statistical power to detect ancestry-associated dependencies among AMR and SAS cell lines and also pointed out limitations to crude continental descriptors. Indeed, dividing cell lines to binary ancestry groups without considering their local ancestry makeup would have resulted in the misclassification of all the admixed cell lines profiled in this analysis. Despite the stark imbalance across all continental ancestry groups, we did maintain sufficient statistical power to detect dependencies associated with cell lines derived from patients of AFR, EAS, or EUR descent [**Figure 1C]**.

We next evaluated whether gene dependencies could be discovered that were significantly positively or negatively associated with a single local ancestry. This analysis revealed 98 such gene dependencies appeared to be associated with either AFR (n = 19), EAS (n = 65), or EUR (n = 56) ancestry; surprisingly, we also detected gene dependencies that are associated with American (n = 2) ancestry, even though we lacked statistical power to detect such associations **[Figure 1D-E]**. Many of these dependency associations (44/98) had a reciprocal relationship with ancestry: each was both positively associated with either EUR or EAS ancestry and negatively associated with the other. This is likely because 91% of the cell lines included in our analysis are either European or East Asian. Interestingly, several ancestry-associated dependencies related to genes related to tyrosine-kinase signaling including *PTPN11* (which encodes SHP2), which is a therapeutic target being tested in clinical trials^13^, and its complex member *GRB2* **[Supplemental Figure 1]**.

We hypothesized that these associations between ancestry and dependencies were due to genetic differences in germline sequences between individuals of different ancestral groups. We therefore searched among SNP loci for dependency qualitative trait loci (d-QTLs) that could explain the differences in dependencies between ancestries. Specifically, we looked genome-wide for the SNP whose genotype was most associated with each dependency. We detected 96 such SNPs across the 98 dependencies. Two of these SNPs (chr9:21340131:A:G and chr9:21338127:A:C) were associated with two different genetic dependencies (IFNE and IFNA16; or IFNA6 and IFNA14), respectively; all four genes are in close proximity on chromosome 9p. Among the 96 SNPs that had the strongest association with dependencies, the genotypes of 35 (36%) were also associated with ancestry **[Figure 2A]**. The SNPs in cis with their dependency gene (< 1 Mb) tended to exhibit more significant associations with dependency than those in trans **[Figure 2B]**. In 41/96 cases the most significant SNPs for each gene were within 1 Mb from the transcription start site of the dependency gene **[Figure 2C]**. Taken together, these data suggest that specific germline variants may dictate dependency on a subset of ancestry-associated genetic dependencies.

**Figure 2:**
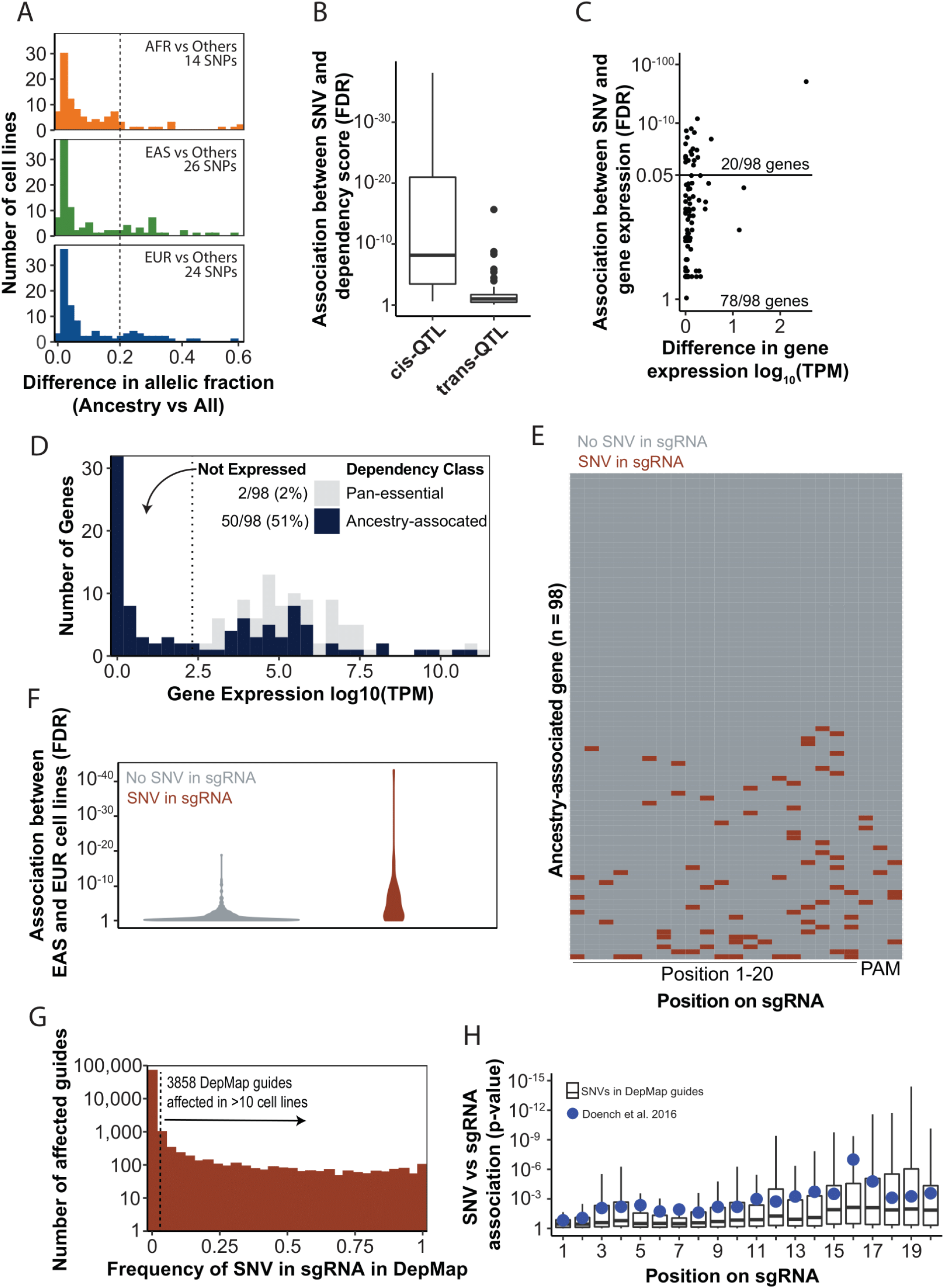
Many apparent ancestry-associated dependencies result from SNP mismatches in sgRNA targeting sequences. **A)** Allele frequency of SNPs that map to sgRNA targeting sequences for ancestry-associated genes. **B)** Significance of the association between the SNP and the gene dependency score for cis and trans d-QTLs. Cis d-QTLs are defined as those within 1 megabase of the transcription start site of the gene in question. **C)** Association between d-QTL SNPs and expression levels of the associated d-QTL gene. **D)** Median expression values for all ancestry-associated dependencies (blue) vs common essential dependencies (grey) across all cell lines profiled in DepMap. **E)** Heatmap indicating loci where SNPs reside on sgRNA target sequences for sgRNAs that target ancestry-associated genes. **F)** Violin plots indicating the distribution of associations between ancestry and gene dependency (as FDR q-values, vertical axis) for dependencies with SNVs in the targeting sequence for at least one sgRNA in at least one cell line (left), vs. all other ancestry-associated dependencies (right). Only dependencies associated with East Asian and European cell lines are shown. **G)** Histogram indicating the number of sgRNAs (vertical axis) against the fraction of Cancer Dependency Map cell lines harboring an SNV in that sgRNA (horizontal axis). **H)** Boxplot showing the association of individual sgRNA depletion scores between cell lines with and without a SNP in the sgRNA targeting sequence (vertical axis) against the location of the SNP in that targeting sequence (horizontal axis). Blue dots indicate the experimentally derived impact of a mismatch in each position from Doench et. al^23^.

We next hypothesized that these d-QTLs conferred differences in gene dependency through their effects on expression of the dependency gene. Across the Cancer Dependency Map, gene expression has been observed to be the strongest predictor of gene dependence^14^. Surprisingly, however, the SNP was associated with expression of the dependency gene in only 20/98 (20%) of cases (q < 0.05). Even among these, cell lines with and without the resistant SNP genotype exhibited median differences in expression of only 8.6%. We suspected that even these differences were not biologically meaningful **[Figure 2C]**. In total, these data suggest that on average, d-QTLs are not modulating the expression levels of their associated genes.

Surprisingly, across all ancestry-associated dependencies, approximately 50/98 (52%) had expression levels below five reads per million, indicating that these genes are weakly expressed or not expressed in a majority of the Cancer Dependency Map cell lines **[Figure 2D]**. The finding that so many genes that appeared to underlie ancestry-associated genetic dependencies were only weakly expressed suggested that the variations in response to CRISPR/Cas9 targeting of many of these genes might reflect something other than true biological differences in gene dependency, such as a technical artifact.

Indeed, differences in cell line response to these sgRNAs might be due to differences in the efficiency with which these sgRNAs were able to induce double strand breaks. The d-QTLs for 47/98 (48%) of ancestry-associated dependencies were in linkage disequilibrium with SNPs in one or more of the sgRNAs targeting the relevant gene **[Figure 2E]**. Mismatches between a CRISPR/Cas9 sgRNA and the target genome preclude guide binding and subsequent genome editing in some circumstances^15,16^, and the frequency of this variation can differ across ancestry groups^17,18^. CRISPR/Cas9-mediated double strand breaks negatively impact cell viability, and can lead to cell death independent of the genomic locus that is targeted by Cas9^19–22^.

These observations support the hypothesis that variation between CRISPR/Cas9 guide and target sequences may explain a substantial fraction of putative ancestry-associated dependency predictions. To comprehensively assess this, we deconstructed the consensus gene dependency scores, which aggregate signals across multiple sgRNAs, into 407 individual sgRNA scores across the 98 dependencies. We then tested the hypothesis that germline SNPs in targeting sequences influenced the differential effects between ancestry groups. As expected, we found that differences in sgRNA depletion between EAS and EUR cell lines was greater for guides with SNPs than for guides without SNPs **[Figure 2F]**. Indeed, this association extended past ancestry-associated variants. Across all sgRNAs in the Cancer Dependency Map, 5.3% have a SNV in their targeting sequence in at least one cell line and 4.3% have such a variant in at least ten cell lines **[Figure 2G]**. Among the latter, which target a total of 2779 genes, 42% of the guides with a SNV in their targeting sequence show a significant association between the presence of a variant and guide dependency. These guides account for 2.2% of all sgRNAs in DepMap **[Figure 2H]**.

Single nucleotide mismatches in an sgRNA targeting sequence can prevent guide binding and the cutting activity of Cas9, with some positions on the sgRNA being less tolerant to mismatches than others^23^. In particular, mismatches in the sgRNA targeting sequence that are closer to the protospacer adjacent motif (PAM) are less tolerant to mismatches than those that are further. We therefore hypothesized that SNV mismatches in sgRNA targeting sequences should impact guide dependency as a function of their distance from the PAM. To test this hypothesis, we compared the location of each SNV mismatch to the magnitude of the difference in dependency between cell lines with and without it. Indeed, mismatches closer to the PAM had a greater impact than those that were further. For example, mismatches in the position farthest from the PAM were not associated with guide dependence (p = 0.15), whereas mismatches in the position closest to the PAM were strongly associated with guide dependence (p < 0.001) **[Figure 2H]**. The impact of mismatches on guide dependence are also significantly correlated with the known impact of mismatches on guide cutting activity^23^ **[Supplemental Figure 2]**.

We found ancestry bias in CRISPR guide design across all ancestry groups, cell models, and CRISPR guide libraries that we evaluated. However, without explicitly accounting for ancestry effects, individuals of predominantly African descent are most affected because people of recent African descent are the most genetically diverse of any continent^24^. This is exemplified when sgRNAs are designed without considering human germline variation. To model this, we chose a random autosomal set of 1,000,000 loci with a canonical NGG PAM site and corresponding protospacer. We limited the selection of these genomic loci to only those that are in protein-coding exons, since most CRISPR-based experiments target coding regions and genomic variability is lower in coding regions than non-coding regions. We then mapped SNPs from 4120 individuals in the individual-level gnomAD dataset to these regions. We found that 62.3% of these sgRNAs contained a SNP in at least one individual, and a median of 1.80% of guides were affected in each individual **[Figure 3A]**. Individuals of African descent, however, were most affected by this artifact (2.17% in AFR, vs 1.78% in all other ancestry groups).

**Figure 3:**
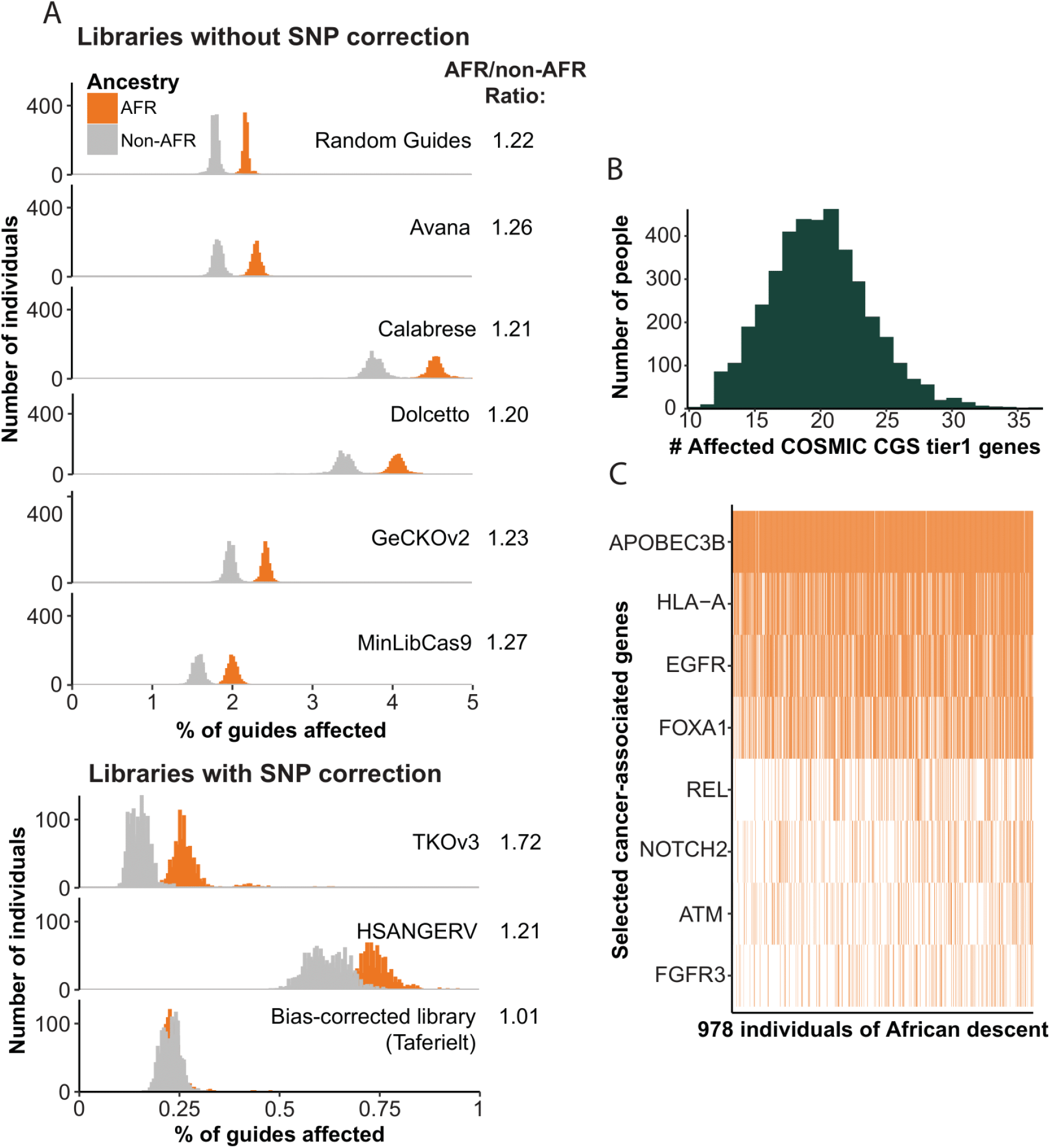
Individuals of African descent are more affected by sgRNA-targeting sequence mismatches than other ancestry groups. **A)** Histograms indicating the frequency (x-axis) which SNPs map to the targeting sequences of guides across 8 CRISPR libraries. Samples are divided into those of African ancestry (orange) and those of other ancestry groups (grey). **B)** Histogram indicating the number of individuals in gnomAD (vertical axis) against the number of COSMIC Cancer Gene Census genes for which that individual’s genome differs from the sgRNA targeting sequence. **C)** The frequency at which sgRNAs from the Avana library are affected by this artifact (orange bars) for 8 selected cancer-associated genes across 978 individuals of African descent.

Multiple factors need to be optimized during the CRISPR sgRNA design process, including maximizing the likelihood that the guide will introduce the intended cut, minimizing the likelihood that the guide will introduce non-specific cuts at additional genomic loci, and minimizing the likelihood of mismatch between the sgRNA and its target due to human variation. For this latter factor, we submit that differences in variant frequencies across populations should be accounted for, to ensure equal efficacy across populations.

Indeed, accounting for human variation in sgRNA design without explicitly accounting for differences in variation across populations does not eliminate the bias against individuals of African ancestry. We mapped germline variants from gnomAD to sgRNA targeting sequences from six genome-scale CRISPR libraries (Avana^23^, Calabrese^25^, Dolcetto^25^, GeCKOv2^26,27^, MinLibCas9^28^, TKOv3^29^, and HSANGERV^30^) **[Figure 3A]**. Among these seven libraries, five (Avana, Calabrese, Dolcetto, GeCKOv2, and MinLibCas9) were designed without attempting to avoid SNP loci in sgRNA targeting sequences. The other two (TKOv3 and HSANGERV) excluded sgRNAs targeting loci with a SNP listed in the db38 and Ensembl 1000 genomes databases, respectively. As might be expected, the five libraries that did not account for SNP variants had the greatest fraction of guides that failed due to human variation, especially in individuals of African descent **[Figure 3A]**. However, all libraries had higher failure rates in African individuals compared to other ancestry groups. Indeed, the ratio between failure rates in African individuals vs other populations was surprisingly constant across all six libraries, ranging from 1.21-1.71. The absolute failure rate was highest in the Calabrese and Dolcetto CRISPRi libraries, likely because these guides map to non-coding regions of the genome.

Although the absolute number of affected CRISPR guides in each individual is small (0.06-4.97% across the six libraries), the impact of this artifact on target discovery in cancer may be large. In the Avana library, for example, 10-36 genes in the COSMIC Cancer Gene Census^31^, are affected by this artifact in each individual **[Figure 3B]**. Many such genes, including EGFR, NOTCH2, ATM, and FGFR3, play important roles in cancer including as oncogenes or tumor suppressors **[Figure 3C, Supplemental Figure 3]**.

Accounting for ancestry-associated human genetic variation can lead to large changes in CRISPR library design. We sought to design an ancestry-agnostic CRISPR library (herein referred to as “Taferielt”) by specifically avoiding regions with high variability in African populations. First, we leveraged the CRISPR sgRNA design tool CRISPick^23,25^ to provide a ranked ordering of sgRNAs with the highest expected cutting rates for each gene while minimizing off-target effects. We then selected the four best sgRNAs for each gene that excluded SNPs with high frequencies in all populations or specifically in African populations (see Methods). We found that these were the four CRISPick top-ranked sgRNAs for only 2,222 (11.5%) of genes. This process resulted in similar rates of mismatches in African individuals (median 0.23%) as in individuals of other ancestry groups (also 0.23%) **[Figure 3A]**.

We also developed new analytical methods to correct for this artifact in existing data. Specifically, we re-calibrated the CRISPR screening data in DepMap to reduce the impact of sgRNA mismatches on the gene-level dependency scores. The corrected version of this dataset was included in the 22Q2 DepMap release.

This artifact does not just affect large-scale CRISPR libraries; rather, it affects all CRISPR-based experiments. We have therefore also developed a web-based tool (www.ancestrygarden.org) that facilitates the discovery of sgRNA sequences that have high mismatch rates across ancestry groups both for the CRISPR libraries profiled in this study and for custom user-input sgRNA sequences.

While we use computational methods that derive continental ancestry groupings to highlight the importance of using diverse reference genomes for developing molecular tools, such continental labels can unintentionally conflate problematic uses of race and human genetic variation. A “multidimensional and continuous conceptualization of ancestry” can resolve some of these issues^32,33^. However, there is a lack of consensus on optimal ways to describe and visualize human genetic variation that are both precise and prevent harm to all people, including groups that have been negatively impacted by racist categorizations.

Herein, we highlight a critical flaw in current CRISPR guide design practices, and we demonstrate the impact this artifact has on discovery of genetic dependencies in cancer. Earlier work also found that genetic variation within CRISPR/Cas9 sgRNA targeting sequences^17^, particularly at therapeutically relevant loci^18^, impacts sgRNA binding. However, until recently it was not possible to systematically understand the impact of this artifact on cancer target discovery. Furthermore, although the potential for this artifact was described over five years ago, most sgRNA design platforms still do not correct for it. We hope to raise awareness of this issue and proposed and implemented a series of solutions to mitigating it.

We previously found that ancestry-associated artifacts can frequently arise in descriptive genomic data^34^; here we find that this also extends to functional genomic data. These findings highlight how widespread such ancestry-associated artifacts are across cancer research, often in ways that are invisible to researchers. The causes of cancer disparities are complex and multifactorial, but biases in basic and pre-clinical research can form an important component. If we hope to make cancer outcomes equitable it is imperative that all forms of ancestry bias are eliminated from cancer research.

---

**Supplemental Figure 1:**
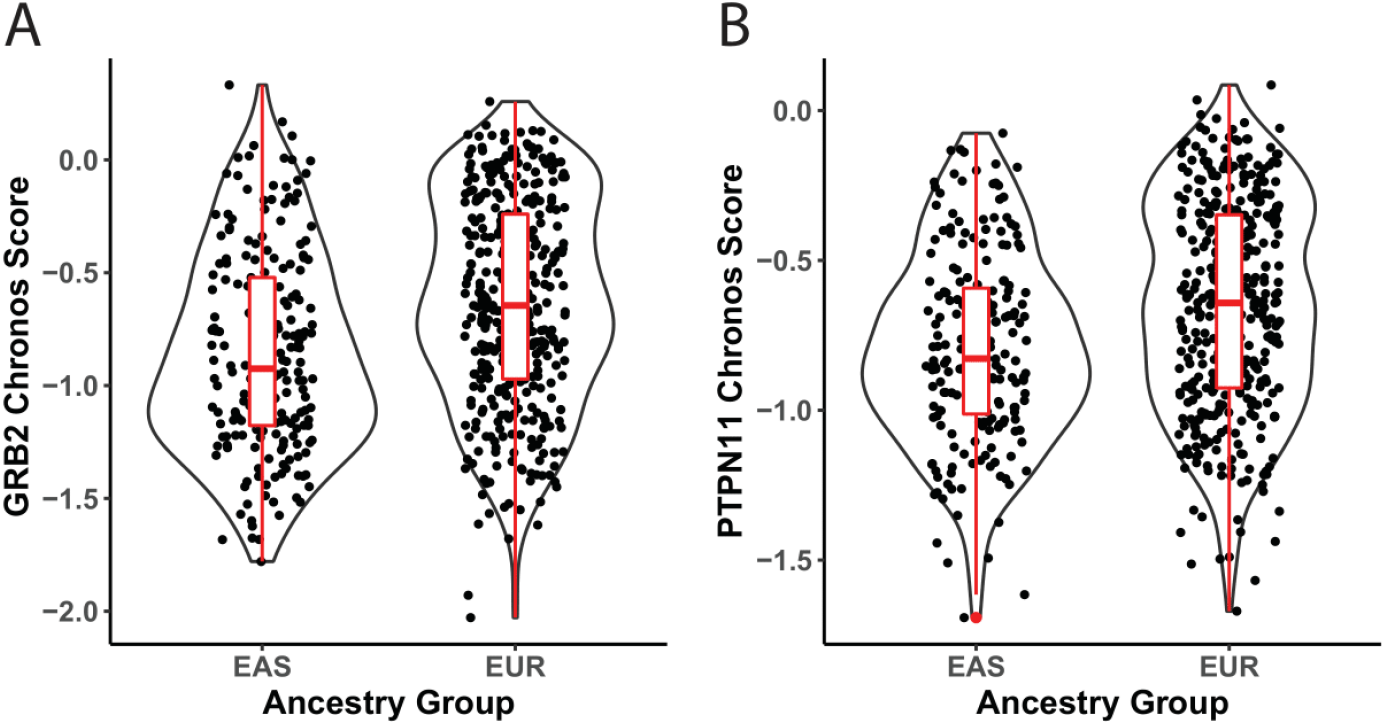
Comparison of GRB2 and PTPN11 dependence in European and East Asian cell lines. Chronos gene dependency scores for **A)** GRB2 and **B)** PTPN11 were compared between European and East Asian cell lines.

**Supplemental Figure 2:**
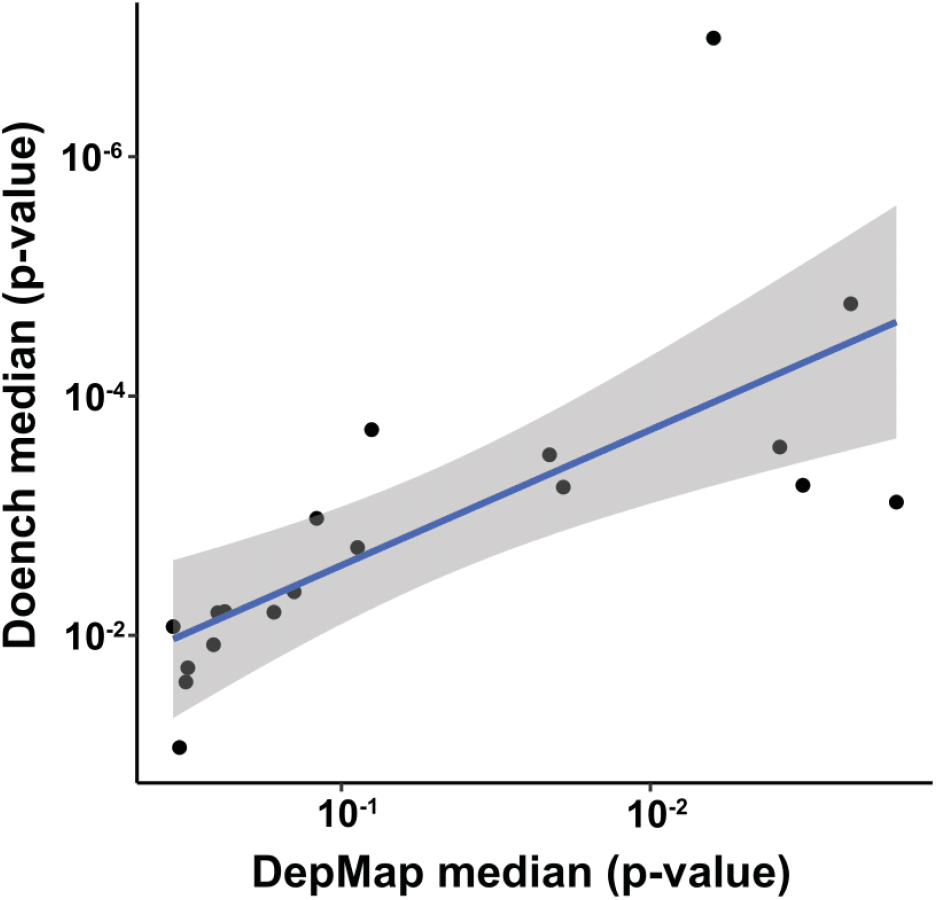
Impact of SNV position on sgRNA cutting activity. Comparison between the impact of SNV mismatches from Doench et al.^23^ (y-axis) with the impact of SNV mismatches across all genes in DepMap (x-axis).

**Supplemental Figure 3:**
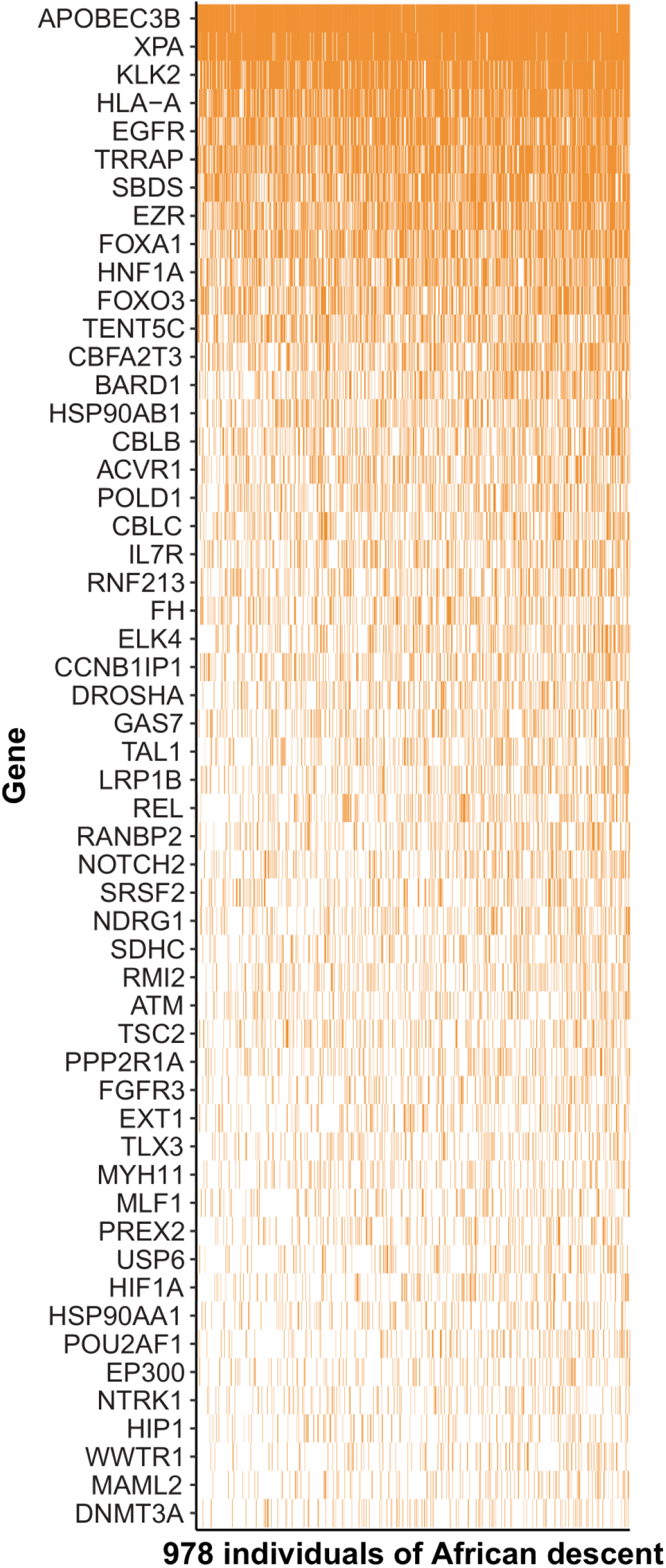
Impact of sgRNA mismatches on individuals of recent African descent. The frequency at which sgRNAs from the Avana library are affected by this artifact (orange bars) in at least 10% of individuals for all COSMIC tier1 genes.

## Methods

### Data sources

CRISPR gene effect, guide-level sgRNA scores, RNA-seq gene expression datasets, and cell line metadata files (Version 22Q1) were downloaded from the Cancer Dependency Map portal (depmap.org). Both the summarized gnomAD and the individual level (HGDP + 1KG) gnomAD data (v3.1.2) were downloaded from gnomad.broadinstitute.org. Guide targeting sequences for each of the genome-scale CRISPR libraries were downloaded from the addgene website.

### Processing SNP6 genotyping calls

Publicly available SNP6 birdseed files^8^ for 994 CCLE cell lines were converted to VCF files as described here (https://software.broadinstitute.org/cancer/cga/contest_prepare2). The resulting VCF files were merged with bcftools. Genotype calls were phased with Eagle (v2.4) and missing genotypes were imputed with Minimac4 (v1.6.6) using the TOPMed reference panel.

### Local ancestry inference

Local ancestry was inferred for 994 CCLE cell lines with RFMix v2 as previously described^3,34^. A set of 2504 unrelated samples profiled as part of the 1000 genomes project were used as a reference panel. The CCLE and 1000 genomes samples were filtered to an intersecting set of variants. RFMix was run with a minimum window size of 0.2 cM.

### Identification of ancestry-associated genetic dependencies

For each gene, cell lines were first binned by their ancestry assignment at the transcription start site of the gene in question. The significance of the association for each ancestry group was computed with a Wilcoxon test comparing cell lines of one ancestry group against cell lines of all other ancestry groups and correcting for multiple comparisons using the False Discovery Rate method^35^.

### Mapping germline variants to sgRNA targeting sequences

We aligned all sgRNA targeting sequences to GRCh38 and discarded those that did not map uniquely to only one genomic loci. SNPs within each individual sample profiled in the HGDP + 1KG gnomAD callset were mapped to each individual guide and guides with a mismatch in at least one allele were identified **(Supplemental Table 2)**.

### Correcting for ancestry bias in The Cancer Dependency Map

We first identified all mismatches between the targeting sequences of guides in the Avana library and the genomic sequences in each individual cell line. Guides with mismatches were excluded only for cell lines with a mismatch when calculating the gene-level dependency (Chronos) score. While this method will reduce the impact of mismatches on false negatives in CRISPR screens, one caveat is that the rate of impacted guides is higher in cell lines with AFR genetic ancestry. This results in a higher rate of eliminated guides in cell lines with AFR genetic ancestry than in cell lines from other genetic ancestry groups.

### Designing an ancestry-agnostic CRISPR library

The top ten sgRNAs for each gene were computed using the CRISPick guide design tool (portals.broadinstitute.org/gpp/public). We then attempted to identify four sgRNAs where there are no mismatches across all samples profiled in gnomAD. For genes lacking four such guides, we selected the guides with the lowest mismatch rates. We imposed the additional restriction that the mismatch frequency within guides may not be present at greater than a 2.5 times rate in AFR individuals than in non-AFR individuals. The identified guide-gene groups were compiled into an ancestry-agnostic “*Taferielt*” CRISPR library.

